# Overexpression of the sugar transporter *VvHT5* turns grapevine into a better host for *Botrytis cinerea*

**DOI:** 10.1101/2025.11.17.687010

**Authors:** Benoit Monnereau, Clément Cuello, Cécile Gaillard, Vincent Lebeurre, Pierre Videau, Olivier Zekri, Pierre Coutos-Thevenot, Sylvain La Camera

**Affiliations:** Écologie et Biologie des Interactions (EBI), UMR7267 Université de Poitiers CNRS, F-86000 Poitiers, France; Novatech, Mercier Groupe, Le Champ des Noels, F-85770 Le-Gué-de-Velluire, France

**Author notes:** These authors contributed equally (Benoit Monnereau, Clément Cuello). Corresponding author: Sylvain La Camera, Écologie et Biologie des Interactions, UMR7267 Université de Poitiers CNRS, F-86000 Poitiers, France.

**Keywords:** Hexose transport, VvHT5, STP13-like transporter, necrotrophic pathogen, grapevine susceptibility, carbon partitioning, *Botrytis cinerea*, tolerance, susceptibility

## Abstract

Sugars are key modulators of plant–pathogen interactions, serving both as metabolic resources and as signaling molecules that influence defense outcomes. In grapevine (*Vitis vinifera*), the hexose transporter VvHT5, an ortholog of *Arabidopsis thaliana* STP13, is strongly induced during fungal infections, suggesting a role in defense-related sugar partitioning. Here, we examined the function of VvHT5 in grapevine susceptibility to the necrotrophic fungus *Botrytis cinerea*. Transgenic lines overexpressing *VvHT5* were generated in two cultivars, Thompson Seedless and Chardonnay, and displayed increased sugar uptake in leaves. Upon *B. cinerea* inoculation, these lines exhibited enhanced lesion development and fungal proliferation, contrasting with the tolerance phenotype previously observed in *A. thaliana* expressing *VvHT5*. Dual RNA-seq analysis revealed that *VvHT5* overexpression triggers a transcriptional landscape favoring carbohydrate metabolism, cell wall remodeling, and attenuation of defense responses, paralleled by upregulation of fungal genes involved in sugar acquisition and virulence. These results demonstrate that while VvHT5 contributes to restricting sugar availability to pathogens in *Arabidopsis*, its constitutive activation in grapevine enhances host suitability for *B. cinerea*. Our findings highlight the complexity of sugar-mediated interactions and underscore the importance of host context in determining the outcome of plant–pathogen competition for carbon resources.

**HIGHLIGHTS:**

- VvHT5 overexpression increases *Botrytis cinerea* susceptibility in grapevine
- Dual RNA-seq uncovers coordinated transcriptional shifts in host and pathogen
- VvHT5 reprograms sugar partitioning and defense-associated metabolism
- Contrasting STP13-like responses between grapevine and Arabidopsis
- Sugar uptake activity of VvHT5 promotes fungal access to host carbon

## 1. INTRODUCTION

Grapevine (*Vitis vinifera* L.) is one of the world’s most economically important fruit crops. Its cultivation is severely threatened by various pathogens with contrasting trophic lifestyles. Among them, *Plasmopara viticola* and *Erysiphe necator*, the causal agents of downy and powdery mildew, respectively, and the necrotrophic fungus *Botrytis cinerea*, responsible for gray mold, are major sources of yield and quality losses (Adrian et al., 2024). *B. cinerea* infects aerial organs such as inflorescences, berries, and leaves, compromising both grape production and quality (Fillinger and Elad, 2016).

*B. cinerea* is a ubiquitous necrotrophic fungus with a broad host range, infecting more than 1,000 vascular plant species across over 500 genera (Elad et al., 2016). Infection begins with the deposition of conidia on plant surfaces and progresses through three main stages of host colonization (Bi et al., 2023). During the early phase, germ tubes differentiate into multicellular appressoria, or infection cushions (Choquer et al., 2021), whose formation is governed by signaling pathways involving G proteins, MAP kinases, adenylyl cyclases, and Ca^2^□/calcineurin-dependent cascades (Beris et al., 2021; Gronover et al., 2001; Harren et al., 2012; Héloir et al., 2019; Klimpel et al., 2002; Zheng et al., 2000). Penetration through these structures marks the onset of the intermediate phase, when the fungus invades host tissues and induces necrotic lesions (Eizner et al., 2017). The late phase is characterized by the rapid expansion of these necrotic areas, culminating in extensive tissue decay.

The ability of *B. cinerea* to successfully infect host tissues relies on a sophisticated arsenal of virulence factors (Choquer et al., 2007; González et al., 2016). Among them, the secretion of plant cell wall–degrading enzymes (PCWDEs) into the apoplasm enables the fungus to dismantle host cell walls and facilitate tissue penetration. Most PCWDEs target carbohydrate components of the plant cell wall and belong to the carbohydrate-active enzymes (CAZymes) family (Blanco-Ulate et al., 2014). These include cellulases that hydrolyze cellulose (Li et al., 2020), xylanases acting on hemicellulose (Brito et al., 2006; Gronover et al., 2004), and pectin-degrading enzymes such as endopolygalacturonases (Leisen et al., 2022; Poinssot et al., 2003), whose activity is facilitated by pectin methylesterases (Kars et al., 2005; Valette-Collet et al., 2003). In addition, *B. cinerea* secretes proteases that degrade structural proteins of the host cell wall (ten Have et al., 2010, 2004).

Among the virulence factors produced by *B. cinerea*, some directly induce host cell death, such as the cell death–inducing proteins (CDIPs) (Bi et al., 2023). For example, purified BcNEP1 and BcNEP2 are sufficient to trigger necrosis in several dicot species including tobacco, broad bean and *Arabidopsis thaliana* (Schouten et al., 2008). Phytotoxins also contribute to virulence, notably Botrydial, the first toxin isolated from *B. cinerea*, whose biosynthesis involves the cytochrome P450 monooxygenase BcBOT1 (Rebordinos et al., 1996; Siewers et al., 2005). Moreover, oxidative burst generated by reactive oxygen species (ROS) promote host cell death, as shown by the reduced virulence of a *B. cinerea* mutant lacking the superoxide dismutase BcSOD1, which is involved in H_2_O_2_ production (Rolke et al., 2004). Finally, *B. cinerea* secretes small RNAs processed by the Dicer-like proteins BcDCL1 and BcDCL2, which can suppress host immunity-related gene expression (Wang et al., 2016; Weiberg et al., 2013), thereby subverting plant defense and enhancing fungal virulence (Veloso and Kan, 2018).

Recognition of the pathogen enables the plant to activate defense mechanisms that restrict infection. These responses depend on the nature of the perceived signals and the receptors involved. Pattern Recognition Receptors (PRRs), localized at the plasma membrane, perceive Pathogen-Associated Molecular Patterns (PAMPs) and/or Damage-Associated Molecular Patterns (DAMPs) from the apoplast (Dodds et al., 2024). Several PRRs have been identified in grapevine (Héloir et al., 2019). Among them, LysM-containing Receptor-Like Kinases (LysM-RLKs) have been the most extensively studied for their role in perceiving chitin and related oligosaccharides, thereby activating defense responses. VvLYK1-1, VvLYK1-2, and VvLYK5-1 have been functionally characterized and shown to restore chitin-triggered responses when expressed in *A. thaliana* mutants deficient in chitin perception (Brulé et al., 2019; Roudaire et al., 2023).

Recognition of PAMPs and DAMPs triggers PAMP-triggered immunity (PTI), initiating a broad range of defense responses (Hudson et al., 2024). Grapevine defense against *B. cinerea* involves complex hormonal signaling networks. Transcriptomic analyses have revealed extensive modulation in expression of genes associated with jasmonic acid, ethylene, salicylic acid and auxin pathways (Agudelo-Romero et al., 2015; Coelho et al., 2019). Downstream of pathogen perception, the regulation of ROS production and detoxification are essential for limiting infection and preventing oxidative damage. Comparative transcriptomic analyses have revealed that tolerant cultivars, including *Vitis amurensis* and *V. vinifera* cv. Syrah, maintain more effective ROS homeostasis and exhibit stronger antioxidant defenses than susceptible cultivars such as *V. vinifera* cv. Red Globe and Trincadeira (Soares et al., 2022; Wan et al., 2021). Defense activation is also accompanied by extensive transcriptional reprogramming of specialized metabolism, leading to the *de novo* synthesis of antimicrobial compounds, known as phytoalexins. In grapevine, many phytoalexins are primarily synthesized *via* the phenylpropanoid pathway, which is initiated by phenylalanine ammonia-lyase (PAL) catalyzing the conversion of phenylalanine into cinnamate (Vannozzi et al., 2012). Upon *B. cinerea* infection, expression of *VvPAL* and several Stilbene Synthases (*VvSTSs*) is induced (Kelloniemi et al., 2015), leading to the accumulation of resveratrol, a stilbene that inhibits conidial germination and fungal development (Adrian et al., 1997; Adrian and Jeandet, 2012; Coutos-Thévenot et al., 2001). In parallel, pathogen perception induces the expression of genes encoding pathogenesis-related (PR) proteins, with antimicrobial activities (Enoki and Suzuki, 2016; Islam et al., 2023). *VvPR2* gene, encoding for β-1,3-glucanase, is strongly induced upon infection by *B. cinerea* or *P. viticola* (Bézier et al., 2002; Mestre et al., 2016). VvPR14 protein, belonging to the lipid transfer protein (LTP) family, is also rapidly induced by *B. cinerea* MAMPs such as ergosterol (Gomès et al., 2003). Moreover, exogenous application of VvLTP4, together with jasmonic acid, prior to infection, increases grapevine tolerance to *B. cinerea* (Girault et al., 2008). Autophagy has also been implicated in grapevine defense against *B. cinerea*, as infection triggers the transcriptional regulation of autophagy-related genes (ATGs) (Zhou et al., 2025). Consistently, overexpression of *VvATG18a* confers greater tolerance to *B. cinerea*, whereas its silencing increases susceptibility.

During pathogen infection, host sugar fluxes are substantially reconfigured through the coordinated induction of cell wall invertases (cwINVs), which hydrolyze sucrose into hexoses, and sugar transporters such as sugar transport protein (STPs), sucrose transporters (SUTs), and sugar will eventually be exported transporters (SWEETs)(Chen et al., 2024; Fotopoulos et al., 2003; Hayes et al., 2010; Veillet et al., 2017). In *A. thaliana, AtSTP13*, encoding a plasma membrane-localized H^+^/hexose symporter, is strongly induced in response to *B. cinerea* and *Pseudomonas syringae* (Lemonnier et al., 2014; Norholm et al., 2006; Yamada et al., 2016). Furthermore, AtSTP13 activity is enhanced following perception of bacterial PAMP, flagellin (flg22), *via* phosphorylation mediated by the co-receptor AtBAK1 (Yamada et al., 2016). Induction of *AtSTP13* orthologs, termed *STP13-like*, by pathogens and PAMPs has been reported in several species, including barley, wheat and *Medicago truncatula*, highlighting a conserved role during PTI (M. Gupta et al., 2021; Huai et al., 2020; Skoppek et al., 2022). Functional studies indicate that STP13-like transporters, which act as plasma membrane-H^+^/hexose symporters, can influence interaction outcomes, functioning either as tolerance or susceptibility factors depending on the host-pathogen combination (Liu et al., 2022). A first approximation suggests that STPs, particularly STP13-like members, contribute to plant tolerance against necrotrophic pathogens, while potentially increasing susceptibility to biotrophic pathogens (Liu et al., 2022). Necrotrophic pathogens preferentially uptake hexoses from the extracellular environment (Doehlemann et al., 2005; Dulermo et al., 2009; Veillet et al., 2017, 2016). STP13-like activity is thought to import apoplastic sugars into the cytosol, thereby restricting sugar availability to necrotrophs and/or supporting host metabolic activity for defense responses. In contrast, biotrophic pathogens form an extra-haustorial matrix that diverts sugars from the host cytosol for their own benefit (Mapuranga et al., 2022). In this context, STP13-like activity may inadvertently increase sugar availability to biotrophic pathogens, thereby contributing to plant susceptibility. In *A. thaliana*, overexpression of *AtSTP13* enhances basal resistance to *B. cinerea*, whereas the *stp13* knockout mutant exhibits increased susceptibility (Lemonnier et al., 2014). Silencing of *TaSTP13* in wheat increases tolerance to *Puccinia striiformis*, and its heterologous overexpression in *A. thaliana* promotes susceptibility to *Golovinomyces cichoracearum* (Huai et al., 2020). In contrast, in *M. truncatula*, silencing of *MtSTP13*.*1* increases susceptibility to *Erysiphe pisi*, whereas transient overexpression in pea induces defense-related gene expression and enhances tolerance (M. Gupta et al., 2021). These contrasting outcomes suggest that STP13-like transporters play interaction-dependent roles, likely by modulating sugar availability at the plant-pathogen interface and contributing to defense signaling. Moreover, the natural Lr67res allele of *TaTP13*, carrying two amino acid substitutions (Arg144Gly and Leu387Val), confers durable resistance in wheat against *P. striiformis, Puccinia graminis* and *Blumeria graminis*, despite its loss of sugar transport activity (Moore et al., 2015). Similar resistance phenotypes were observed when *Lr67res*-like mutations were introduced into *HvSTP13* from barley and *MtSTP13*.*1* from *M. truncatula* (M. Gupta et al., 2021; Milne et al., 2019; Skoppek et al., 2022). Recent evidence suggests that this resistance is independent of hexose transport, potentially involving anion flux-related mechanisms (Milne et al., 2024, 2023).

Interestingly, sugar fluxes are dynamically reprogrammed not only in the host but also in the pathogen. Analyses of *B. cinerea* sugar transporter genes, particularly hexose transporters, reveal distinct regulation patterns during infection of sunflower cotyledons and *A. thaliana* leaves, highlighting intense competition for sugars between host and pathogen (Dulermo et al., 2009; Veillet et al., 2017, 2016). Functional characterization of *BcSDR1*, an essential gene for full pathogenicity, further links glucose transport to fungal virulence, underscoring the critical role of sugar acquisition in disease progression (Si et al., 2022; Zang et al., 2018).

In grapevine, expression of the grapevine hexose transporter *VvHT5*, the ortholog of *AtSTP13*, is strongly induced in leaves upon infection by multiple fungal pathogens, including *E. necator, P. viticola, Eutypa lata*, and *B. cinerea* (Cardot et al., 2019; Hayes et al., 2010; Monnereau et al., 2025). This induction is accompanied by increased expression of *VvcwINV*, suggesting coordinated control of apoplastic sugar fluxes. *VvHT5* may also act in concert with SWEET transporters, including *VvSWEET4* in leaves and *VvSWEET7* in berries, both of which are induced by *B. cinerea* (Breia et al., 2020; Chong et al., 2014; Monnereau et al., 2025). SWEETs play a key role in modulating plant tolerance or susceptibility, particularly those localized at the plasma membrane that facilitate sugar efflux between the cytosol and the apoplast (Chong et al., 2014; P. K. Gupta et al., 2021; Liu et al., 2022). Pathogens, including bacteria and fungi, exploit SWEET transporters to increase apoplastic sugar availability by promoting the export of sucrose and hexoses from host cells.

To dissect the functional significance of *VvHT5*, we previously examined its ectopic expression in *A. thaliana*, where it enhanced tolerance by restricting necrosis spread during the late phase of infection (Monnereau et al., 2025). In the present study, we extend this analysis to grapevine. Stable transgenic lines overexpressing *VvHT5*, exhibiting increased sugar uptake in leaves, were generated in two cultivars. Surprisingly, these lines displayed enhanced susceptibility to *B. cinerea*. Dual RNA-seq analyses revealed that *VvHT5* overexpression reshapes the dialogue between grapevine and *B. cinerea*, establishing sugar transport as a decisive player in plant-pathogen interactions, tipping the balance between resistance and susceptibility.

## 2. MATERIAL AND METHODS

### 2.1. Plant materials and growth conditions

#### 2.1.1. Induction of embryogenic calli

Ovaries were collected from immature flowers of *V. vinifera* cv Chardonnay (Cha) clone 124 grown in the field at Vix (France) and placed on induction medium to initiate internal embryogenic calli. The medium consisted of Murashige and Skoog (MS) basal salts, supplemented with 30 g·L□^1^ sucrose, adjusted to pH 5.8, and supplemented with 4.52 µM 2,4-dichlorophenoxyacetic acid (2,4-D), 1.23 µM indole-3-butyric acid (IBA), and 1.34 µM naphthaleneacetic acid (NAA). Explants were subcultured at monthly intervals, and the emergence of embryogenic calli was monitored from the sixth month onwards. *V. vinifera* cv Thompson Seedless (TS) explants were obtained under the same conditions.

Embryogenic cultures were maintained on GM+NOA medium as described by Coutos-Thévenot et al. (1992), using liquid medium for TS embryogenic cells and solid medium for Cha embryogenic calli.

#### 2.1.2. Plant growth conditions

Grapevine plants of TS and Cha cultivars, including both wild-type and transgenic lines, were grown in a greenhouse in a 3:1 mixture of BP-Substrat 264 (Klasmann-Deilmann GmbH) and vermiculite. Plants were maintained under a long-day photoperiod (16 h light / 8 h dark) with a minimum temperature of 22 °C during the day and 18 °C at night.

### 2.2. VvHT5 cloning and grapevine transformation

The full-length coding sequence of *VvHT5* (1,611 bp) was cloned into the pENTR-D-TOPO vector (Invitrogen) as described by Monnereau et al. (2025) and subsequently transferred into the pK7WGF2 binary vector (Karimi et al., 2002) via LR recombination. The *VvHT5* coding sequence was fused downstream of the GFP coding sequence and placed under the control of the Cauliflower mosaic virus 35S (CaMV 35S) promoter. The resulting construct was introduced into *Agrobacterium tumefaciens* strain GV3101 (pMP90) by electroporation.

Grapevine transformation was performed following the method described by Mauro et al. (1995), with minor modifications. TS embryogenic cells and Cha embryogenic calli were co-cultured with *A. tumefaciens* (OD□□□ = 0.2, pretreated with 100 µM acetosyringone) in liquid GM + β-naphthoxyacetic acid (NOA) medium under stirring for 30 minutes at 28°C. Cells and calli were then spread onto sterile Whatman paper placed on a Petri dish containing solid GM + NOA medium. The Petri dish was kept in the dark at 26°C for 48 hours.

TS cells were washed and transferred to liquid GM^0^ medium (GM without NOA), for embryogenesis (Coutos-Thevenot et al., 1992), supplemented with cefotaxime (200 µg.mL^−1^) to eliminate residual *A. tumefaciens* and paromomycin (2 µg.mL^−1^) to select transformed cells, as described by Mauro et al. (1995). The medium was refreshed regularly over several weeks. Embryos were transferred to McCown Woody Plant medium (Duchefa), supplemented with 20 g.L^−1^ sucrose, pH 5.8, and 3.5 g.L^−1^ gelrite, along with cefotaxime (200 µg.mL^−1^) and paromomycin (2 µg.mL^−1^).

Cha calli were washed and transferred first to liquid GM + NOA medium, supplemented with cefotaxime (200 µg.mL^−1^). For selection and regeneration, calli were then transferred to McCown Woody Plant medium (Duchefa), supplemented with 20 g.L^−1^ sucrose, 0.89 µM 6-benzylaminopurine, 0.01 µM indole-3-butyric acid, pH 5.8, and 3.5 g.L^−1^ gelrite, along with cefotaxime (200 µg.mL^−1^) and kanamycin (50 µg.mL^−1^).

Resulted plantlets were propagated by cuttings and multiplied on a standard grapevine propagation medium, consisting of McCown Woody Plant medium (Duchefa) with 30 g.L^−1^ sucrose, pH 5.8, and 7.5 g.L^−1^ type-A agar (Sigma).

Transgene genomic integration was verified by PCR using a primer pair designed to amplify the T-DNA region spanning the CaMV 35S promoter and the T35S terminator, including the GFP-*VvHT5* fusion sequence (Supplementary Table S1). PCR was performed on genomic DNA extracted from leaves of *in vitro* plantlets using the NucleoSpin Plant II extraction kit (Macherey-Nagel).

### 2.3. Western blot analysis

Microsomal fractions were prepared from mature grapevine leaves as described by Lemonnier et al. (2014). Membrane proteins were separated by SDS-PAGE on a 10% polyacrylamide gel and transferred to a nitrocellulose membrane (Hybond ECL, GE Healthcare) for western blot analysis. The membrane was incubated with either anti-GFP antibodies (Euromedex-GeneTex) or anti-VvHT5 antibodies (Monnereau et al., 2025). Antibody detection was performed using the ECL Plus Western Blotting Detection Kit (GE Healthcare) and visualized with the Amersham Imagen 600 (GE Healthcare).

### 2.4. Glucose uptake and soluble sugar quantification

Radiolabeled glucose uptake into leaf discs was performed as previously described by Monnereau et al. (2025).

Soluble sugars were extracted from grapevine leaves (150 mg) following the method described by Veillet et al. (2017). Briefly, frozen leaf powder was thawed and homogenized in 1.5 mL of MCE extraction buffer (60% methanol, 25% chloroform, 15% water). After centrifugation at 1,200 g for 10 minutes at 20°C, the supernatant was collected. The remaining sugars in the pellet were extracted twice with 600 µl of MCE. The supernatants were pooled and mixed with 0.6 volumes of water. After centrifugation at 1,200 g for 15 minutes at 20°C, the upper aqueous phase was collected and evaporated using a Speed Vac (miVac QUATTRO concentrator). The pellet was resuspended in water. Sugars extracted from grapevine samples were further purified using Sep Pak C18 columns (Waters). Soluble sugars (sucrose, glucose and fructose) were quantified using the Sucrose/D-Fructose/D-Glucose Assay kit (Megazyme) according to the manufacturer instructions.

### 2.5. Culture of *Botrytis cinerea*, infection methods and disease scoring

*B. cinerea* B05.10 strain (Staats & van Kan, 2012) was grown from a glycerol stock of conidia on solid Difco Potato Dextrose Agar (PDA) medium (Becton-Dickinson) at 22°C under long-day conditions until sporulation. Conidia were harvested by suspending spores in sterile water followed by filtration through Miracloth paper (Merck).

#### 2.5.1. Assessment of susceptibility to *B. cinerea*

Susceptibility to *B. cinerea* was evaluated on leaf discs (2 cm diameter) excised from mature grapevine leaves. Each disc was placed on Whatman 3 MM CHR paper (Cytiva) saturated with sterile water in a 120 × 120 mm Petri dish (Corning). A 6 µL drop of *B. cinerea* B05.10 conidial suspension (5 × 10□ conidia.mL□^1^), prepared in one-quarter diluted Difco Potato Dextrose Broth (PDB; Becton-Dickinson), was applied to the center of the abaxial surface of each disc. Petri dishes were incubated in a water-saturated mini-greenhouse, in the dark, for 72 h. Necrotic areas were quantified using an IMAGING-PAM M-series chlorophyll fluorometer (WALZ). Leaf discs were irradiated with actinic light (100 µmol·m□^2^·s□^1^) for 500 s, followed by a pulse of saturating light. Image analysis was performed based on the quantum yield of non-regulated energy dissipation [Y(NO)], which allows optimal discrimination between necrotic (non-photosynthetic) and healthy (photosynthetic) tissues, using ImageJ software (Schneider et al., 2012). Necrotic areas were measured using the MicroArray Profile plugin (OptiNav). Necrotic areas, expressed as the percentage of total disc surface, were compared among genotypes using a Kruskal-Wallis non-parametric test, followed by Dunn’s post hoc multiple comparison test with Benjamini-Hochberg correction for false discovery rate (dunnTest function implemented in the FSA package v.0.10.0; Ogle et al., 2015). Groups that did not differ significantly (α = 0.05) were assigned the same letter using the cldList function from the rcompanion package (v.2.5.0; Mangiafico, 2025). Lesions were further categorized into three size classes following Lemonnier et al. (2014): small lesions were defined as smaller than the first quartile of lesion sizes observed in the corresponding wild type (WT), medium lesions fell within the interquartile range of the WT, and large lesions exceeded the third quartile of the WT. Lesion size distributions among classes were compared to their respective wild-type controls using pairwise Chi-square tests.

#### 2.5.2. Whole-leaf infection assay for RNAseq analysis

For whole leaf infection, leaves were placed in water-saturated mini-greenhouses and the abaxial surface was sprayed with a *B. cinerea* B05.10 conidia suspension (10^5^ conidia.mL^−1^) prepared in 1/4 diluted PDB. The mock treatment was carried out by spraying the leaves with 1/4 PDB solution, without conidia. Mini-greenhouses were incubated in the dark in a growth chamber (21°C, 60% relative humidity). Leaves were collected and frozen in liquid nitrogen 48 hours post post-inoculation (hpi). Infected samples included three biological replicates (rep1 to rep3).

### 2.6. *B. cinerea* quantification in infected leaves

Genomic DNA (gDNA) was extracted using the NucleoSpin Plant II kit (Macherey-Nagel) from mature healthy grapevine leaves, grapevine leaves infected with *B. cinerea* (48 hpi), and *B. cinerea* mycelium grown on PDA solid medium. The gDNA extracted from healthy leaves and fungal mycelium was serially diluted and quantified using a Nanodrop 1000 spectrophotometer (Thermo Scientific).

Quantitative PCR (qPCR) was performed using primer pairs specific to *VvPDS* for grapevine and *BcCUTA* for *B. cinerea* (Gachon and Saindrenan, 2004) (Supplementary Table S1) to generate two standard curves relating Ct values to the amount of grapevine or fungal gDNA. gDNA from infected leaves was amplified with both primer pairs, and the corresponding Ct values were used to determine the relative amounts of plant and fungal DNA in each sample based on the standard curves. Three biological replicates were analyzed for TS lines and two for Cha lines.

### 2.7. Fluorescent imaging

The GFP signal in transgenic grapevine leaves was initially confirmed using a fluorescence stereomicroscope (MacroView MVX10, Olympus). For higher resolution at the subcellular level, images were acquired by confocal microscopy (Olympus FV3000). Excitation was performed with laser lines at 488 nm for GFP and 640 nm for chlorophyll, and fluorescence emission was detected in the 500–540 nm and 650–750 nm ranges, respectively.

### 2.8. Gene expression analysis

#### 2.8.1. RNA extraction

Total RNA extraction from grapevine leaves was performed using the CTAB extraction method described by Chang et al. (1993) and modified as described by Morin et al. (2022). Briefly, 100 mg of frozen leaf powder was thawed and homogenized in 1 mL of CTAB buffer (Tris/HCL 100 mM, pH 8, CTAB 2%, EDTA 25mM, NaCl 2M, spermidine 3.44 mM). RNA was subsequently purified by two successive extractions with chloroform/isoamyl alcohol (24:1). After RNA precipitation with isopropanol, purification is conducted on a Qiagen column (RNeasy Mini Kit).

#### 2.8.2 Gene expression analysis by quantitative real-time PCR (qRT-PCR)

RNA extracted from grapevine leaves was treated with DNase (Sigma) and reverse transcribed into cDNA using M-MLV Reverse Transcriptase (Promega). qRT-PCR was carried out on the Light Cycler 480 thermocycler using the GoTaq qPCR Master Mix reagent (Promega). Gene expression was normalized to the expression of the reference gene *Glyceraldehyde 3-phosphate dehydrogenase* (*VvGAPDH)*, using the 2-ΔCt method (Schmittgen and Livak, 2008). Primers were selected from literature and are listed in Supplementary Table S1.

#### 2.8.3. RNA sequencing

For the non-treated condition, whole mature leaves were harvested from acclimated plants and immediately frozen in liquid nitrogen. Five biological replicates (rep1 to rep5) were collected. For *B. cinerea* and mock treatments, three biological replicates were included (rep1 to rep3).

RNAseq analysis was performed using RNA extracted from TS transgenic (TS#14) and wild-type TS. Prior sequencing, RNA was precipitated with lithium chloride. RNAseq library preparation and sequencing was performed by GENEWIZ platform (Azenta Life Sciences) following their *in-lab* procedure. Strand-specific RNAseq libraries were prepared with polyA selection and sequenced on an Illumina NovaSeq 6,000 system in paired-end 2 x 150 bp mode. A range of 56 to 89 million raw reads was obtained per library. Quality control and contaminant detection, including adapters and primers, are performed using FastQC (v0.11.9) (Andrews, 2010). Trimming and filtering are performed using Fastp (v0.25.0) (Chen et al., 2018), with a length of at least 50 bp and a Phred score of 30 or higher. Raw RNA-seq data have been deposited under the BioProject accession numbers PRJNA1363860.

#### 2.8.4. Differential expression and functional enrichment analysis

RNAseq reads (54 to 85 million quality-filtered RNAseq reads per RNAseq library) were mapped to a concatenated genome composed of telomere-to-telomere *Vitis vinifera* cv ‘Pinot Noir’ genome (T2T_PN40024, Shi et al., 2023) and *B. cinerea* B05.10 genome (ASM14353v4 [GenBank ID: GCA_000143535.4], van Kan et al., 2017) using STAR (v2.7.11b, Dobin et al., 2013) with the following parameters: SortedByCoordinate, max number of mismatches normalized to the mapped lengths: 0.05, outFilterIntronMotifs: RemoveNoncanonicalUnannotated. Expression quantification was determined by FeatureCounts (v2.1.1, Liao et al., 2014). The counting matrix (Supplementary Table S2) was subsequently divided into two matrices, one for *Vitis* genes, the other for *Botrytis* genes. Non-expressed genes were removed using remove_nonexp function implemented in BioNERO package (v1.14.0, Almeida-Silva & Venancio, 2022) with the following options: method = “percentage”, min_exp = 10, min_percentage_samples 3/22. RNAseq data were normalized using TMM implemented in the edgeR package (v4.4.2, Robinson et al., 2010). Gene-wise RNAseq count were then analysed using a negative binomial generalized linear model for these different analyses: (i) *Vitis* genes differential expression was assessed between TS wild-type and TS#14 non-treated conditions with line main effect, (ii) *Vitis* genes differential expression was line-independently assessed between Mock-inoculated and *B. cinerea* (Bc)-infected conditions with infection status as main effect, (iii) *Botrytis* genes differential expression was assessed between TS wild-type and TS#14 Bc-infected conditions with line main effect. A Benjamini-Hochberg adjusted *p*-value of 0.05 and a log-FoldChange of |2| was used as the cut-off criterion to identify significantly differentially expressed genes.

Functional annotations were downloaded from grapedia (https://grapedia.org) for *V. vinifera* T2T_PN40024 and from NCBI for *B. cinerea* ASM14353v4 (BioProject: PRJNA15632). Gene Ontology (GO) enrichment analysis is performed for DEG lists using topGO (v2.58.0, Alexa et al., 2022). Overrepresented GO terms are identified using a two-sided Fisher’s Exact test, with significance defined as a Benjamini-Hochberg adjusted *p*-value ≤ 0.05.

### 2.9. Statistics and graphs

Principal Component Analyses (PCA) were performed using the mixOmics R package (v6.30.0, Rohart et al., 2017). Heatmaps of differentially expressed genes were generated with the ComplexHeatmap package (v2.22.0, Gu, 2022). All graphical visualizations, including bar plots, Venn diagrams, PCA, Heatmaps and enrichment plots, were produced using the ggplot2 package (v3.5.2, Wickham, 2011) and ggpubr package (v0.6.0, Kassambara, 2022) in the R environment (v 4.4.1, R Core Team, 2024), or using GraphPad Prism version 10.6.1 (GraphPad Software, La Jolla California USA).

## 3. RESULTS AND DISCUSSION

### 3.1. Generation of VvHT5-overexpressing grapevine lines through genetic transformation

To investigate the role of VvHT5 in regulating sugar-mediated defense in grapevine, we generated stable transgenic plants that constitutively overexpress this STP13-like hexose transporter (Monnereau et al., 2025). The *VvHT5* coding sequence from *Vitis vinifera* cv Chardonnay was subcloned into the pK7WGF2 vector (Karimi et al., 2002) as an N-terminal GFP–VvHT5 fusion under the control of the CaMV 35S promoter. Because grapevine is difficult to transform and regenerate (Campos et al., 2021), a GFP tag was fused to *VvHT5* to facilitate the visualization of transformed tissues and to monitor both transformation efficiency and transgene expression. Genetic transformation was performed using *Agrobacterium tumefaciens* strain GV3101 on embryogenic cells derived from TS and Cha. These cultivars are widely used for *Agrobacterium*-mediated grapevine transformation and have been successfully transformed with strain GV3101 in previous studies (Campos et al., 2021; Capriotti et al., 2023; Dai et al., 2015; Fan et al., 2008; Iocco et al., 2001; Su et al., 2018; Wang et al., 2017).

After antibiotic selection *in vitro*, 38 TS and 23 Cha kanamycin-resistant plantlets were recovered. Several lines failed to survive *in vitro* culture and the subsequent steps (leading to multiplication and acclimation). Some lines exhibited irregular PCR amplification patterns or abnormal morphology. Ultimately, two TS and six Cha transgenic lines were successfully acclimated to greenhouse conditions by transferring to soil in pots, together with their respective WT controls (Supplementary Figure S1a).

The number of surviving transformants was consistent with previously reported efficiencies for *Agrobacterium*-mediated transformation of grapevine, which remains challenging by low transformation and regeneration rates (Campos et al., 2021; Iocco et al., 2001).

### 3.2. Characterization of transgenic lines

To assess transgene expression in the eight transgenic lines, *VvHT5* transcript levels were quantified in healthy mature leaves of acclimated plants by qRT-PCR (Figure 1a). Since the transgene encodes a GFP fusion, we aimed to distinguish transgene expression from endogenous *VvHT5*. To this end, separate GFP- and VvHT5-specific primers were used to monitor transgene expression and total *VvHT5* expression (Supplementary table S1), the latter including the endogenous *VvHT5* (Supplementary Figure S1b).

**Figure 1.**
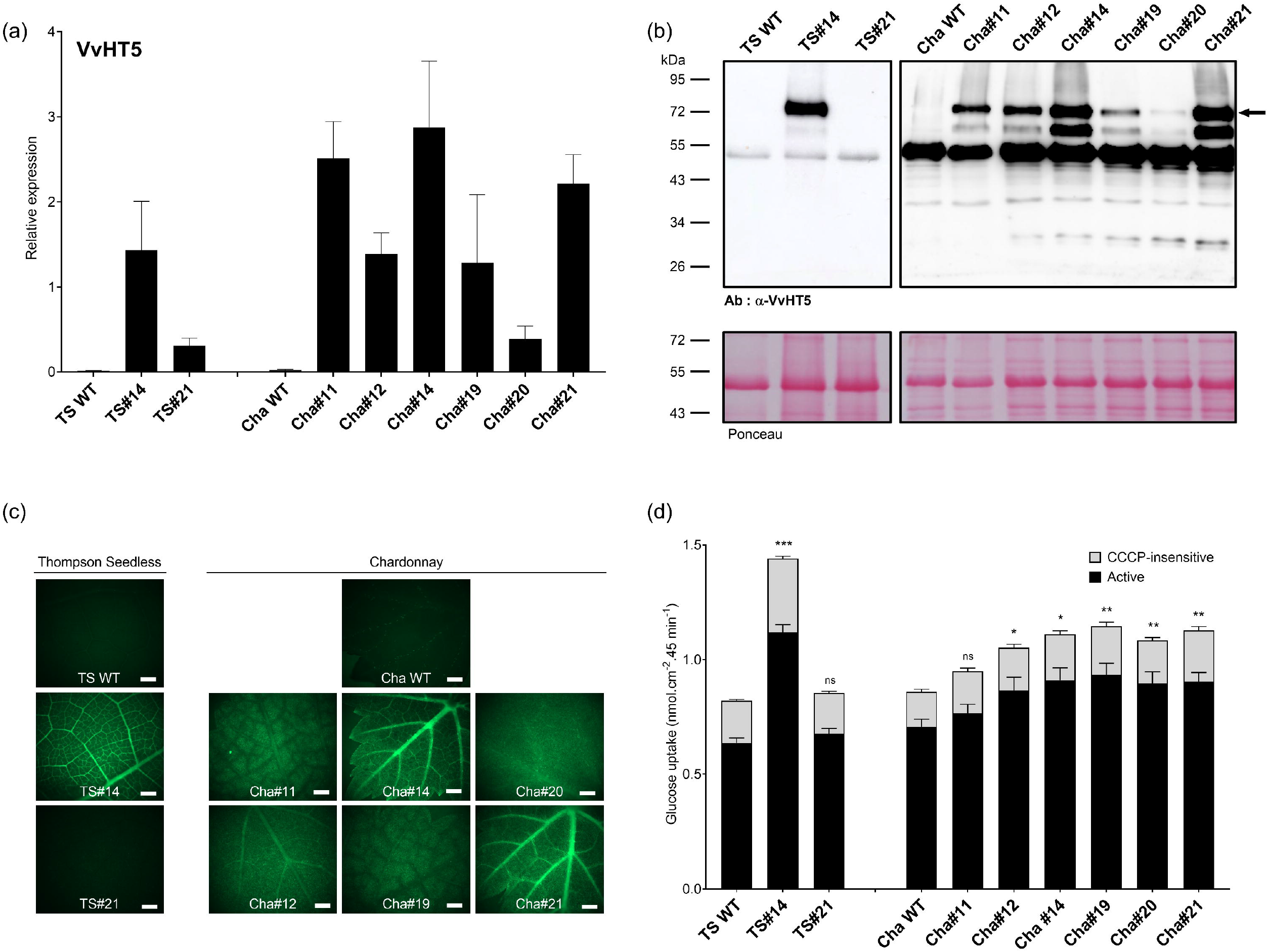
Characterization of *VvHT5-*overexpressing *Vitis* lines. (a) *VvHT5* expression levels in leaves of wild-type and transgenic TS and Cha lines. Gene expression was quantified by qRT-PCR from healthy mature leaves and normalized to the reference gene *VvGAPDH*. Data represent mean ± SD (three biological replicates). (b) Accumulation of the GFP–VvHT5 protein in transgenic TS and Cha lines assessed by Western blot. Microsomal fractions extracted from healthy mature leaves were probed with anti-VvHT5 antibodies. The black arrow indicates the VvHT5-specific band. (c) GFP fluorescence in leaves of wild-type and *VvHT5*-overexpressing TS and Cha lines. GFP signal emitted by the GFP–VvHT5 protein was detected using a stereoscope (scale bar: 1 mm). (d) Glucose uptake in healthy mature leaf discs of wild-type and *VvHT5*-overexpressing TS and Cha lines. Uptake of [^1^□C]-labeled glucose was measured in the presence or absence of the protonophore CCCP. Data represent mean ± SEM (three biological replicates, six technical replicates each). Asterisks indicate significant differences relative to the corresponding wild-type according to pairwise Wilcoxon tests (alternative = “less”) using Benjamini-Hochberg adjustment method (\**p* < 0.05, \*\**p* < 0.01, \*\*\**p* < 0.001).

Although *VvHT5* expression levels varied among the lines, all transgenic lines exhibited enhanced and constitutive expression, whereas transcript levels in WT plants remained low and barely detectable. Among the transgenic lines, TS#14 exhibited the highest overexpression (~130-fold), Cha#11, Cha#14, and Cha#21 showed strong overexpression (~100-fold), Cha#12 and Cha#19 displayed moderate overexpression (~50-fold), and lower levels were observed in TS#21 and Cha#20. The identical expression patterns observed using the GFP-specific primers indicate that the enhanced expression in the transgenic lines is attributable to transgene expression rather than changes in endogenous *VvHT5* expression.

A polyclonal antibody against VvHT5 (Monnereau et al., 2025) was used to assess protein accumulation in microsomal fractions from leaves of transgenic plants (Figure 1b). No VvHT5 protein was detected in the corresponding WT plants, consistent with its low constitutive expression. All lines, except TS#21, showed substantial VvHT5 accumulation, which correlated with transcript levels (Figure 1a). TS#14, Cha#14, and Cha#21 exhibited particularly high protein accumulation. The absence of detectable protein in TS#21, despite transgene transcripts, highlights potential post-transcriptional or post-translational regulation, a phenomenon commonly observed in transgenic plants. This result was further confirmed using a GFP-specific antibody (Supplementary Figure S1c), which revealed the same protein expression patterns across the transgenic lines, highlighting that the observed accumulation is driven by the transgene. To go further, we examined tissue-specific expression in leaves by monitoring GFP fluorescence using a wide-field fluorescence stereomicroscope (Figure 1c). GFP fluorescence was observed only in the transgenic lines accumulating the fusion protein. While some lines (Cha#11, Cha#12, Cha#19, Cha#20) displayed uniform fluorescence across the leaf blade, the signal was particularly strong in the vascular tissues of TS#14, Cha#14, and Cha#21, which also exhibited the highest protein levels. Confocal imaging of TS#14 leaves further revealed a peripheral signal in epidermal cells (Supplementary Figure S1d), supporting the plasma membrane localization of this H^+^/monosaccharide symporter, as previously reported for VvHT5 (Hayes et al., 2010; Monnereau et al., 2025).

Taken together, these results demonstrate that the enhanced transcript levels of the *GFP– VvHT5* transgene are effectively translated into protein accumulation, which is specifically detectable in the transgenic lines and absent in WT plants. The correlation between transcript and protein levels, along with the tissue-specific GFP fluorescence, confirms the functionality of the fusion construct. Notably, strong accumulation in vascular tissues and peripheral localization in epidermal cells support the predicted plasma membrane targeting of this H^+^/monosaccharide symporter, consistent with its proposed role in sugar transport. These findings provide a solid foundation for further physiological studies, including *in planta* transport assays to evaluate the functional impact of VvHT5 overexpression in grapevine.

In Monnereau et al. (2025), we previously demonstrated the functionality of VvHT5, both with and without an N-terminal GFP fusion, showing that it mediates high-affinity [^14^C]-glucose uptake. Based on this, we assessed [^14^C]-glucose uptake in grapevine leaf discs. Most transgenic lines exhibited significantly higher uptake rates than corresponding WT (Figure 1d). Notably, TS#14 displayed the highest glucose uptake, with a 76% increase relative to WT, while the transgenic Cha lines showed significant increases of up to 34%. TS#21 and Cha#11 showed comparable values. The absence of increased uptake in TS#21 was consistent with the lack of VvHT5 protein accumulation (Figures 1b, c and Supplementary Figure S1c) whereas in Cha#11, it may be due to additional, unidentified factors affecting transporter activity.

Exposure of leaf discs to the protonophore CCCP allowed measurement of passive (CCCP-insensitive) glucose uptake, thereby distinguishing the active component of total glucose uptake, which is mediated by proton-coupled hexose transporters. TS#14 showed the largest increase (+76% relative to WT), while Cha#14, Cha#19, Cha#20, and Cha#21 exhibited significant increases of approximately 30%, and Cha#12 displayed a trend toward increase of 22% (*p* = 0.06458) (Supplementary Table S3). This analysis confirmed that the increased total glucose uptake observed in transgenic lines was accompanied by a statistically significant increase in active glucose uptake, indicating that it results from VvHT5 activity.

The activity of plasma membrane H^+^/hexose transporters, such as VvHT5, enables the uptake of hexoses from the apoplasm into cells. The fate of these imported sugars remains unclear, but they are hypothesized to support increased metabolic needs during pathogen attack or to help regulate intracellular processes (Yamada and Mine, 2024). In the present study, the content of soluble intracellular sugars remained unchanged in TS#14, Cha#19, and Cha#21, which exhibited the strongest increases in VvHT5 expression and glucose uptake, compared to WT (Supplementary Figure S1e). These results are consistent with previous observations in *Arabidopsis* expressing VvHT5 (Monnereau et al., 2025) but contrast with overexpression of TaSTP13, which led to increased glucose accumulation and enhanced susceptibility to powdery mildew (Huai et al., 2020). Considering the diversity of sugar transporters with distinct substrate specificities and subcellular localizations, it is likely that their collective activity offsets the increased sugar influx mediated by VvHT5, thereby preserving intracellular sugar balance in the transgenic grapevine lines. This also raises the possibility that VvHT5 may serve functions beyond hexose transport or be involved in additional, unrelated cellular processes.

Together with previous reports, these findings confirm that VvHT5 shares key features with STP13-like transporters. Members of this clade, including AtSTP13 (Lemonnier et al., 2014; Norholm et al., 2006; Yamada et al., 2016), TaSTP13 (Huai et al., 2020), MtSTP13.1 (M. Gupta et al., 2021), and HvSTP13 (Skoppek et al., 2022), are plasma membrane H^+^/monosaccharide symporters whose expression is induced during pathogen infection. Similarly, VvHT5 transcript levels increase in response to infections by both necrotrophic and biotrophic pathogens, including *B. cinerea, E. lata, E. necator*, and *P. viticola* (Cardot et al., 2019; Hayes et al., 2010; Monnereau et al., 2025), consistent with its classification as a stress-responsive STP13-like transporter.

### 3.3. Overexpression of *VvHT5* enhances grapevine susceptibility to *B. cinerea*

The transgenic grapevine lines generated in this study represent the first reported grapevine plants with altered sugar transporter activity, by elevated VvHT5 expression and increased glucose uptake. This successful challenge of stable genetic transformation in grapevine provides a unique system to investigate whether VvHT5 influences the outcome of the grapevine-*B. cinerea* interaction under controlled conditions.

Leaf discs from VvHT5-overexpressing lines displaying enhanced sugar uptake were drop-inoculated with *B. cinerea* and disease resistance was first assessed by measuring the size of the resulting necrotic lesions. Under our experimental conditions, the TS and Cha cultivars exhibited contrasting susceptibility levels, with TS being more susceptible to *B. cinerea* than the Cha genotype, showing an average necrotic area of 36.3% compared to 22.4% for Cha (Figure 2a). Overall, both transgenic Cha lines and the TS#14 line exhibit significantly higher necrotic areas than their wild-type counterpart, suggesting that the genetic modifications enhanced disease severity to infection. Each lesion was categorized as small, medium, or large, and the distribution of lesion sizes in transgenic lines was compared to that observed in their respective WT (Figure 2b and Supplementary Table S4). All tested transgenic lines showed a significantly altered distribution, with a higher proportion of large lesions and fewer medium and/or small ones, indicating enhanced susceptibility. Importantly, this phenotype was observed in two distinct grapevine cultivars, TS and Cha, which display contrasting basal tolerance levels, thereby reinforcing the robustness of the result. Notably, lines TS#14 and Cha#20 showed a pronounced increase in severe symptoms, accompanied by a marked decrease in less affected areas. To go further, fungal colonization was quantified at 48 hpi by measuring the gDNA ratio (*B. cinerea*/*V. vinifera*) in whole infected detached leaves (Figure 2c). Consistent with its pronounced disease phenotype (Figure 2d), TS#14 showed substantially higher fungal colonization, whereas all other transgenic lines displayed colonization levels comparable to their respective controls at this stage of infection. These results indicate that VvHT5 overexpression increases grapevine susceptibility to *B. cinerea*, particularly in the more susceptible TS genotype, where enhanced VvHT5 expression seems to promote fungal growth. It suggests that VvHT5 may facilitate pathogen development by providing additional hexose resources to the fungus in infected tissues. Interestingly, this phenotype contrasts with the enhanced resistance previously observed in *A. thaliana* upon overexpression of either *VvHT5* or its ortholog *AtSTP13* (Lemonnier et al., 2014; Monnereau et al., 2025). This opposite outcome emphasizes the limitations of the *A. thaliana* model for predicting host responses in grapevine and, more broadly, in crop species with distinct physiological and metabolic characteristics (Bevan et al., 2025; Korwin Krukowski et al., 2020; Piquerez et al., 2014; Uauy et al., 2025). In *A. thaliana*, STP13-like overexpression was proposed to deplete apoplastic sugars, thereby restricting nutrient availability to the necrotrophic pathogen *B. cinerea* (Lemonnier et al., 2014; Monnereau et al., 2025). By contrast, overexpression of TaSTP13 in *A. thaliana* increased susceptibility to the biotrophic pathogen *G. cichoracearum* (Huai et al., 2020), a phenotype associated with higher leaf glucose content, leading the authors to hypothesize that STP13-like transporters could increase cytoplasmic glucose availability and facilitate nutrient access for haustorium-forming pathogens. In grapevine, as in *A. thaliana*, VvHT5 overexpression was not accompanied by detectable changes in leaf soluble sugar content (Supplementary Figure 1e) (Monnereau et al., 2025). Therefore, the enhanced disease severity observed in transgenic grapevine lines infected with the necrotroph *B. cinerea* does not appear to result from altered sugar content, suggesting that other physiological or metabolic factors, potentially host species-specific or related to the pathogen’s lifestyle, modulate the impact of VvHT5 on host–pathogen interactions. Although *B. cinerea* is generally described as a necrotroph, recent studies have highlighted an initial biotrophic phase during early infection, during which the fungus may transiently suppress host defenses and modulate sugar metabolism (Bi et al., 2023; Veloso and Kan, 2018). It is therefore tempting to hypothesize that VvHT5 activity may contribute to this early phase. A premature or excessive induction of VvHT5 could, as reported for TaSTP13 and TaSTP6 in wheat (Huai et al., 2020, 2019), create a favorable environment for the fungus, either supporting increased fungal growth during its biotrophic stage or accelerating the transition to necrotrophy. Such regulatory mechanisms could influence the expression or activity of VvHT5, and merit further investigation to better understand how *B. cinerea* adapts its infection strategy to host physiology

**Figure 2.**
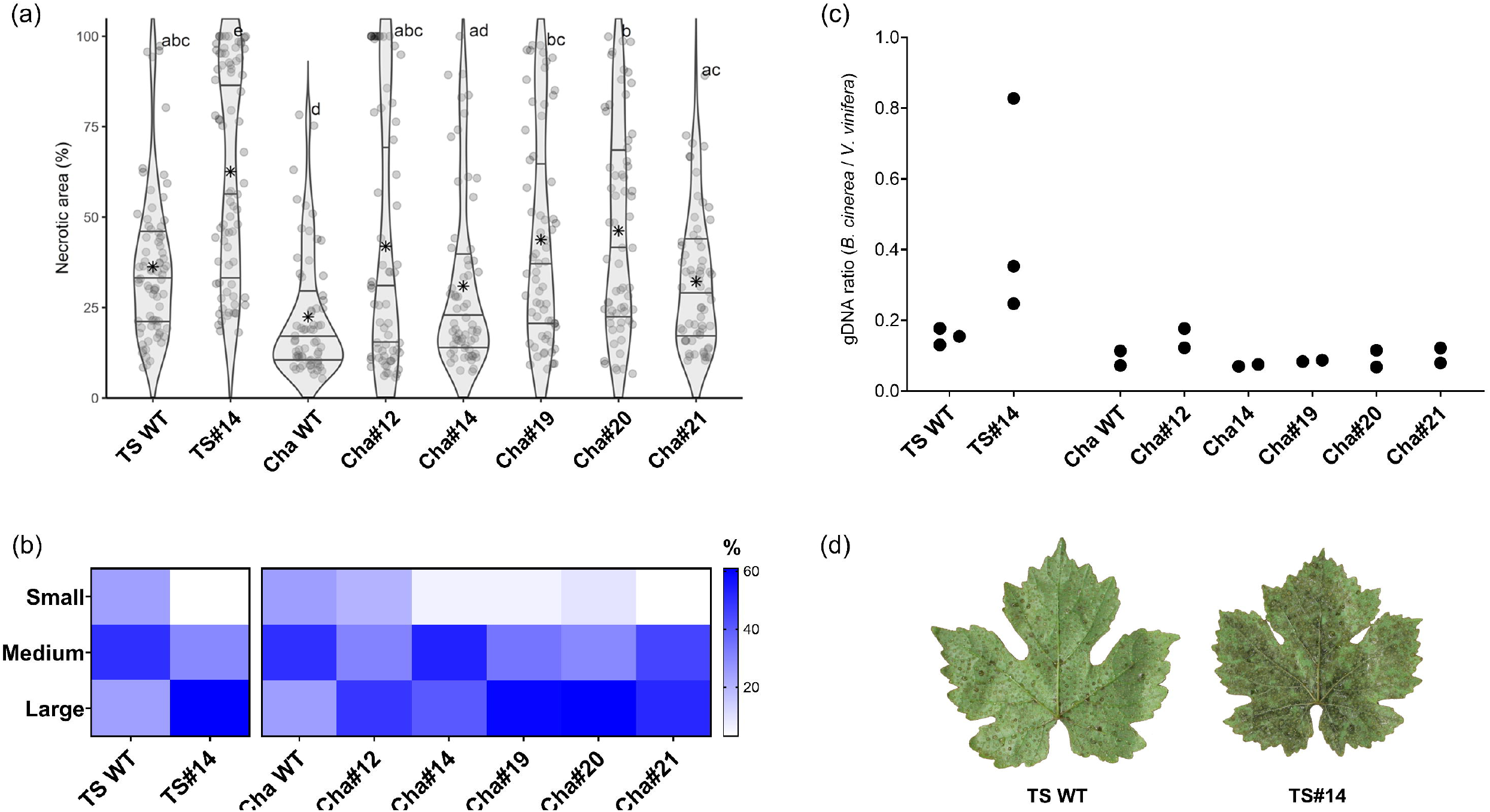
Susceptibility of *VvHT5*-overexpressing *Vitis* lines to *Botrytis cinerea*. (a) Percentage of necrotic area measured on mature leaf discs at 72 h post-inoculation (hpi) with *B. cinerea* for each line. For Thompson Seedless lines, 76 leaf discs were analyzed for TS WT and TS#14; for Chardonnay lines, 64 leaf discs were analyzed per line. Individual observations are shown as semi-transparent jittered points. Violin plots represent lesion size distributions, with internal lines indicating the first, second, and third quartiles, and a black asterisk marking the mean. Letters above violins correspond to groups identified by Dunn’s post hoc test (α = 0.05, Benjamini–Hochberg correction). Lines sharing letters are not significantly different, whereas those without letters in common differ significantly. (b) Foliar symptom severity for all lines at 72 hpi, presented as a heatmap based on a three-class scale. Dark blue indicates the highest percentage (60%), and white indicates the lowest (0%). (c) Relative quantification of *B. cinerea* in infected leaves of wild-type and VvHT5-overexpressing lines. Mature leaves spray-infected with *B. cinerea* were sampled at 48 hpi for DNA extraction. The ratio of *B. cinerea* to grape gDNA (BcCUTA / VvPDS) was determined by qPCR using organism-specific primers and DNA standards. Black dots indicate biological replicates (three for Thompson Seedless lines, two for Chardonnay lines). (d) Symptoms of *B. cinerea* infection on whole detached leaves from TS WT plants and transgenic line TS#14 at 48 hpi.

STP13-like transporters are known to be tightly connected to plant defense signaling. Their expression and activity are induced by PAMPs, such as flg22, chitin and its derivates (M. Gupta et al., 2021; Skoppek et al., 2022; Yamada et al., 2016). Moreover, silencing of *MtSPT13*.*1* in *M. truncatula* reduces the induction of salicylic acid-dependent defense marker genes, and PR proteins during *E. pisi* infection, while its transient overexpression in pea leaves enhances *PR* gene expression before and after infection (M. Gupta et al., 2021). In grapevine, VvHT5 may function as part of a regulatory system linking to the activation of defense pathways, similar to the roles proposed for other STP13-like transporters in diverse plant–pathogen interactions. Accordingly, we hypothesize that the increased susceptibility observed in VvHT5-overexpressing grapevine lines could result from alterations in transcriptional regulation in both the host and the fungus, rather than from changes in sugar availability.

### 3.4. Global transcriptional landscape

To investigate the transcriptional consequences of *VvHT5* overexpression in grapevine and its impact on the interaction with *B. cinerea*, we performed RNAseq on TS wild-type and TS#14 under three conditions: non-treated (NT), mock-inoculated (Mock), and infected with *B. cinerea* (Bc) at 48 hpi. Reads were mapped onto a concatenated reference genome combining *V. vinifera* PN40024 and *B. cinerea* B05.10, which ensured efficient and reliable alignment across all samples (Figure 3a). Although mapping specificity tended to decrease slightly in inoculated conditions, this effect was consistent across genotypes. Notably, the proportion of reads assigned to *B. cinerea* genome increased markedly in Bc-infected TS#14 compared to TS wild-type, in line with the great susceptibility of the transgenic line. These observations are consistent with the quantification of fungal DNA (figure 2b), which also revealed a higher accumulation of *B. cinerea* biomass in the TS#14 line compared to the wild-type.

**Figure 3.**
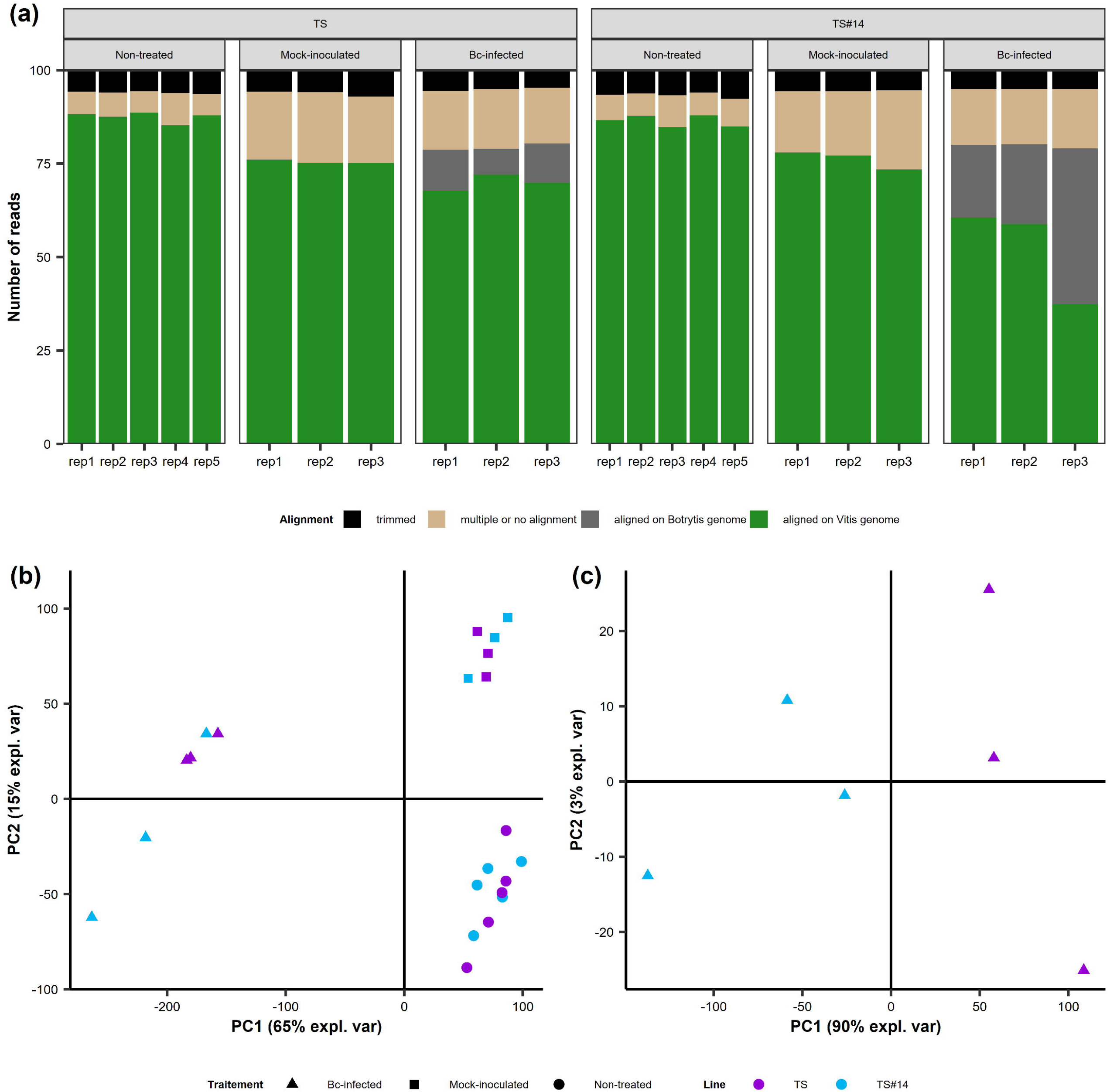
Overview of RNA-seq data quality and transcriptomic variation across conditions. (a) Proportion of reads mapped to the *Vitis vinifera* genome (green), the *Botrytis cinerea* genome (grey), unmapped of multiply aligned reads (tan), or low-quality reads (black). (b-c) Principal Component Analysis of genes expressed in *Vitis* (b) or *B. cinerea* (c) across all sample conditions. *Purple: TS, Blue: TS#14, Triangle: Bc-infected, Square: Mock-inoculated, Circle: Non-treated*.

Principal component analysis (PCA) of grapevine gene expression confirmed the strong reproducibility of biological replicates and showed that non-treated and mock-inoculated samples were largely indistinguishable, irrespective of genotype (Figure 3b). In contrast, infection with *B. cinerea* revealed a clear separation between TS wild-type and TS#14, highlighting genotype-dependent transcriptional responses. A complementary analysis of the fungal gene expression further revealed a striking divergence between the two host backgrounds (Figure 3c), underscoring that *B. cinerea* transcriptional program is profoundly reconfigured by the grapevine genetic background it colonizes. These global patterns pointed to profound transcriptional adjustments in both partners during infection. Yet, they did not reveal why the transgenic line is more vulnerable. To uncover the molecular basis of this heightened susceptibility, we next examined gene expression changes in detail through differential analysis.

### 3.5. Overexpression of *VvHT5* has negligible constitutive effects

To assess the constitutive impact of *VvHT5* overexpression, we compared gene expression profiles between TS wild-type and TS#14 under NT conditions. The analysis revealed only 11 differentially expressed genes (DEG), with seven more highly expressed in TS wild-type and four more highly expressed in TS#14 (Table 1). Although counterintuitive, such a limited basal transcriptomic footprint is consistent with the notion that membrane sugar transporters primarily modulate flux rather than act as transcriptional regulators. In line with this, sugar-transporter transgenic lines often display modest constitutive changes but marked, context-dependent phenotypes upon stress or during development. Consistently, RNAseq analysis of grapevine calli overexpressing *VvSWEET10* showed that transcriptional reprogramming was largely restricted to genes related to sugar transport and metabolism, indicating limited pleiotropic effects under control conditions (Zhang et al., 2019). Similarly, transcriptome profiling of *osstp15* rice highlighted target shifts at the shoot base (*e*.*g*., cytokinin synthesis and signaling linked to altered sugar status) rather than broad constitutive deregulation (Li et al, 2024). Importantly, the minimal basal divergence here allows us to attribute the pronounced differences seen upon infection (Figure 3b-c) to infection-triggered genotype-dependent responses rather than broad constitutive rewiring.

**Table 1.** Constitutively differentially expressed genes between TS wild-type and TS#14. ^*a*^ *log(FoldChange);* ^*b*^ *Benjamini-Hochberg adjusted p-value*

| Gene ID | eggNOG annotation | mean expression |  | logFC | adjusted<br>p-value |
| --- | --- | --- | --- | --- | --- |
|  |  | TS | TS#14 |  |  |
| Vitvi05_01chr05g07090 | Belongs to the sugar transporter (TC 2.A.1.1) family | 568,8 | 70068,4 | -6,98 | 2,4E-91 |
| Vitvi05_01chr18g32880 | NA | 4,2 | 45,8 | -3,47 | 1,7E-03 |
| Vitvi05_01chr18g05690 | NA | 242 | 2591,6 | -3,39 | 1,3E-28 |
| Vitvi05_01chr14g08250 | Reverse transcriptase (RNA-dependent DNA polymerase) | 144 | 1131,2 | -3,00 | 2,1E-18 |
| Vitvi05_01chr11g18790 | Belongs to the UDP-glycosyltransferase family | 1096 | 264,2 | 2,15 | 2,9E-04 |
| Vitvi05_01chr11g18310 | Catalyzes xyloglucan endohydrolysis and/or endotransglycosylation. | 34 | 7,2 | 2,24 | 4,9E-02 |
| Vitvi05_01chr11g18830 | Belongs to the UDP-glycosyltransferase family | 525 | 98,4 | 2,47 | 1,8E-08 |
| Vitvi05_01chr07g14320 | NA | 469,2 | 81 | 2,54 | 6,3E-03 |
| Vitvi05_01chr11g18730 | antiporter activity | 31,4 | 5 | 2,67 | 5,8E-03 |
| Vitvi05_01chr07g16180 | NA | 47,6 | 4 | 3,49 | 1,4E-05 |
| Vitvi05_01chr11g18740 | Belongs to the multi antimicrobial extrusion (MATE) (TC 2.A.66.1) family | 1273,4 | 21 | 5,95 | 2,8E-45 |

Beyond *VvHT5*, which unsurprisingly showed the strongest difference in expression, the other DEG represented a heterogeneous set. They comprised a reverse transcriptase more abundant in TS#14, two UDP-glycosyltransferases (UGT), two putative transporters and a gene related to primary cell wall construction preferentially expressed in TS wild-type. Importantly, none of these genes are directly associated with classical plant determinant of the resistance/susceptibility responses (Adrian et al., 2024), supporting the view that the increased susceptibility of TS#14 to *B. cinerea* does not result from a constitutive predisposition at the transcriptional level. However, changes in UGT expression could still have consequences for specialized metabolism, particularly flavonol/anthocyanin glycosylation, given the well-established roles of UGTs such as VvUFGT in grapevine and UGT78D1/2 in *A. thaliana* in modulating flavonoid conjugate profiles and directing phenylpropanoid fluxes (Boss et al., 1996; Yonekura-Sakakibara et al., 2012; Yonekura-Sakakibara and Hanada, 2011). Likewise, one of the putative transporters (Vitvi05_01chr11g18740) is a member of the multi-antimicrobial extrusion family with recognized function in the compartmentation of flavonoids (Marinova et al., 2007; Pérez-Díaz et al., 2014), implying that modest transcription shifts might alter the availability of defense-related metabolites without overtly reprogramming immunity. Finally, xyloglucan endo-transglycosylase/hydrolase (XTH) remodel xyloglucan and thereby influence cell wall mechanics, a determinant of plant-pathogen outcomes (Bellincampi et al., 2014), and xyloglucanase activity from *B. cinerea* (BcXYG1) can itself act as an elicitor (Zhu et al., 2017), underscoring why altered XTH expression could matter even in the absence of a “defense pathway” signature.

While no single defense pathway emerges from the DEG analysis, subtle adjustments in specialized metabolism, metabolite transport, and cell-wall remodeling remain plausible modulators of the heightened susceptibility observed in TS#14 upon infection. Overall, these results indicate that VvHT5 overexpression exerts minimal influence on grapevine basal transcriptional activity, yet such subtle shifts could shape the physiological context in which infection unfolds. To explore how these latent differences translate into divergent outcomes upon *B. cinerea* challenge, we next examined the transcriptional responses of both genotypes during infection.

### 3.6. Differential transcriptional responses of grapevine genotypes during infection

To disentangle infection-specific responses from experimental effects, we compared gene expression between mock-inoculated and *B. cinerea*-infected samples for each genotype separately. This analysis revealed a massive transcriptional reprogramming, with 9,589 DEGs in TS#14 and 9,096 in TS wild-type (Figure 4a-b). Most of these changes were shared, as illustrated by the 3,070 genes upregulated and the 4,996 downregulated in both genotypes in response to infection. The large overlap indicates that the two lines mount broadly similar responses to *B. cinerea* and suggests that their contrasting susceptibilities are linked to more subtle regulations. Notably, *VvHT5* was part of the shared response (TS#14: *p*-value = 8.64e^−11^, log2FC = 3.44; TS wild-type: *p*-value = 2.93e^−41^, log2FC = 9.16), reaching similar expression levels in TS#14 and TS wild-type (Figure 6b). This similarity suggests that the transcriptional changes and the higher susceptibility of TS#14 may result from signaling events occurring prior to 48 hpi, rather than from differences in transporter abundance at this timepoint. Consistently, time-resolved atlas studies of the *Arabidopsis*-*B. cinerea* interaction show that early (6-24 hpi) transcriptional waves, especially in defense and hormone networks, prefigure disease outcome (Windram et al., 2012).

**Figure 4.**
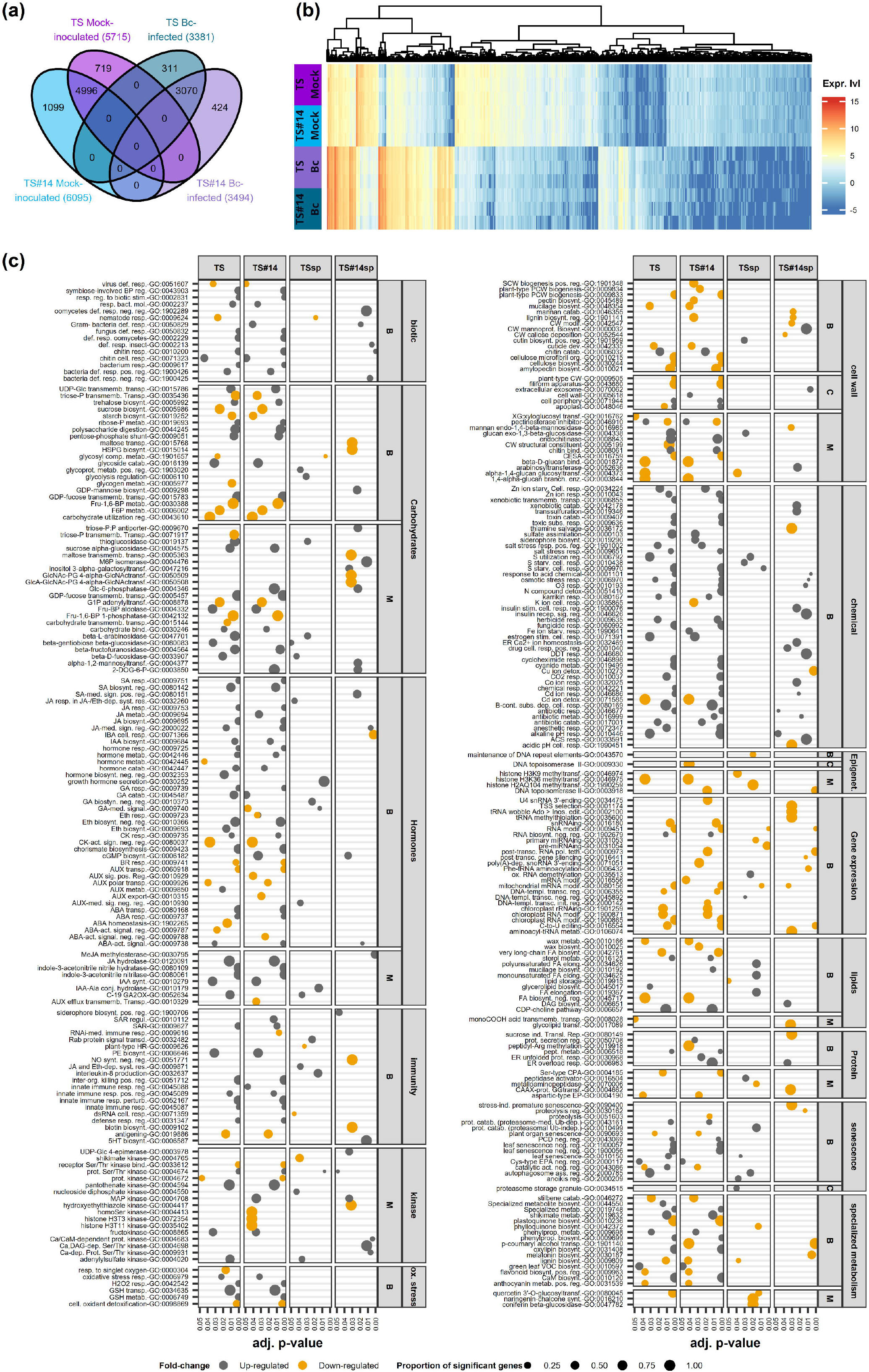
Transcriptomic response of *Vitis vinifera* to *Botrytis cinerea* infection. (a) Venn diagram of upregulated (dark colours) and downregulated (light colours) genes in response to infection TS (purple) and TS#14 (blue). (b) Heatmap of *Vitis* differentially expressed genes (DEGs) in response to *Botrytis cinerea. Purple: TS Mock-inoculated, Blue: TS#14 Mock-inoculated, dark purple: TS Bc-infected, dark blue: TS#14 Bc-infected* (c) Gene Ontology enrichment analysis of *Vitis* DEGs. *B: biological process, C: cellular compartment, M: molecular function. TSsp/TS#14sp: enrichment performed on genes specifically differentially expressed in TS or TS#14, respectively. Grey: GO term significantly over-represented in genes induced by B. cinerea; Yellow: GO term significantly over-represented in genes repressed by B. cinerea*.

To gain insight into these specific mechanisms, we performed a Gene Ontology (GO) enrichment analysis on the sets of DEGs unique to each genotype (Figure 4c, Supplementary Table S5). In TS wild-type, enriched categories pointed to a coordinated activation of defense-related processes, including systemic hormone signaling, particularly jasmonic acid. This hormone signature matches the canonical view that jasmonic acid/ethylene pathways underpin effective defenses against necrotrophs, whereas salicylic acid-dominated states tend to be less protective in these pathosystems (Glazebrook, 2005; Pieterse et al., 2012). The analysis also revealed an increase in lipid and sulfur metabolism, together with the regulation of autophagy. Autophagy components (*e*.*g*., ATG18, ATG7) contribute to resistance to necrotrophic fungi, including *B. cinerea*, by constraining runaway cell death and tuning defense output (Lai et al., 2011; Lenz et al., 2011). Moreover, terms associated with glycoprotein and cutin biosynthesis, as well as the activation of cutinases, were significantly enriched, suggesting reinforcement and remodeling of the plant’s surface barrier. Genetic and biochemical evidence indicates that modifying cuticle composition or permeability profoundly alters *B. cinerea* resistance through barrier function and early defense signalling (Aragón et al., 2021; Bessire et al., 2007). The production of specialized metabolites such as phylloquinone was also promoted consistent with the mobilization of multiple metabolic layers to counter *B. cinerea*. More broadly, specialized metabolites form integral layers of innate immunity against fungal pathogens (Piasecka et al., 2015). Functional terms also indicated a reinforcement of genetic and epigenetic regulation, including repression of microRNA activity, modulation of dsRNA responses and histone methylation. Such reinforcement of RNA- and chromatin-level control aligns with the need to rapidly coordinate large defense gene sets during *B. cinerea* attack in time-series transcriptomes (Windram et al., 2012). Altogether, these enrichments suggest that TS wild-type mount a multifaceted defense strategy aimed to both at restricting *B. cinerea* progression through autophagy and cell wall strengthening and producing metabolites that may directly counteract *B. cinerea*. In addition, enrichment of sulfur- and glutathione-related processes is compatible with improved redox buffering and detoxification of ROS/toxins, both implicated in resistance to *B. cinerea* (Künstler et al., 2020; Zechmann, 2020).

By contrast, TS#14 displayed a markedly different enrichment pattern. Hormonal regulations involved a more complex interplay between jasmonic acid and salicylic acid signaling. Given the well-described antagonism between salicylic acid and jasmonic acid, such co-activation can blunt effective anti-necrotroph responses (Glazebrook, 2005; Pieterse et al., 2012). Several transport-related processes were specifically induced, including calcium signaling, triose-phosphate and phosphoenolpyruvate transport, while others such as maltose, *p*-coumaryl and glycolipid transport, were repressed. This pattern suggests carbon repartitioning that pathogens can exploit, in line with evidence that microbial pathogens manipulate plant sugar transporters to access host carbon (Breia et al., 2021; Liu et al., 2022; Veillet et al., 2017). TS#14 also showed enriched responses to abiotic and biotic cues including drought, bacteria, insects, and oomycetes. At the genetic level, functional categories pointed to repression of post-transcriptional gene silencing (PTGS), sucrose-induced translational regulation, and post-translational modification. Notably, *B. cinerea* has been shown to deploy small RNA effectors that alter immunity (Cheng et al., 2025; Wang et al., 2016; Weiberg et al., 2013). In this light, the altered PTGS signature in TS#14 may reflect changes in small RNA-mediated regulation during infection, but the directionality and causal link cannot be inferred from GO terms alone. In parallel, senescence, lignin and polysaccharide remodeling and cell wall reinforcement appeared inhibited, whereas mannoprotein accumulation, mannan, xenobiotic catabolism, and antibiotic biosynthesis were favored. Such broad stress- and detoxification-skewed signatures, rather than targeted antifungal programs, are commonly observed in susceptible *Botrytis*-host interactions. Comparative RNAseq and time-series studies report that susceptibility correlates with heightened ROS/redox and cell-death modules, whereas more resistant genotypes mount earlier, better-coordinated jasmonate/ethylene, cell-wall and antioxidant responses (Haile et al., 2020, 2017; Wan et al., 2021; Windram et al., 2012; Yang et al., 2020).

Together, these results indicate that while TS wild-type mobilizes multiple layers of defense, including hormone signaling and chemical/physical barriers, to slow down *B. cinerea*, TS#14 displays a more diffuse and seemingly maladaptive response. The latter is characterized by transcriptional signatures that suggest the pathogen overwhelms host defenses, redirecting host metabolism rather than encountering an effective counterattack.

### 3.7. Transcriptional reprogramming of *B. cinerea* in response to *VvHT5* overexpression

We next compared *B. cinerea* gene expression profile during infection of the two hosts. A total of 286 genes were differentially expressed, of which 265 showed higher expression on TS#14 host and only 21 on TS wild-type host (Figure 5a). This striking asymmetry indicates that overexpression of *VvHT5* in the host triggers a major transcriptional reprogramming in the pathogen. Such a skew is consistent with the well-documented induction of fungal CAZymes and sugar transport systems when readily usable carbon becomes available *in planta* (Blanco-Ulate et al., 2014; Lacrampe et al., 2021; Reboledo et al., 2021), raising the possibility that VvHT5-driven apoplastic carbon dynamics relax nutrient limitation and broadly activate *B. cinerea*-acquisition programs.

**Figure 5.**
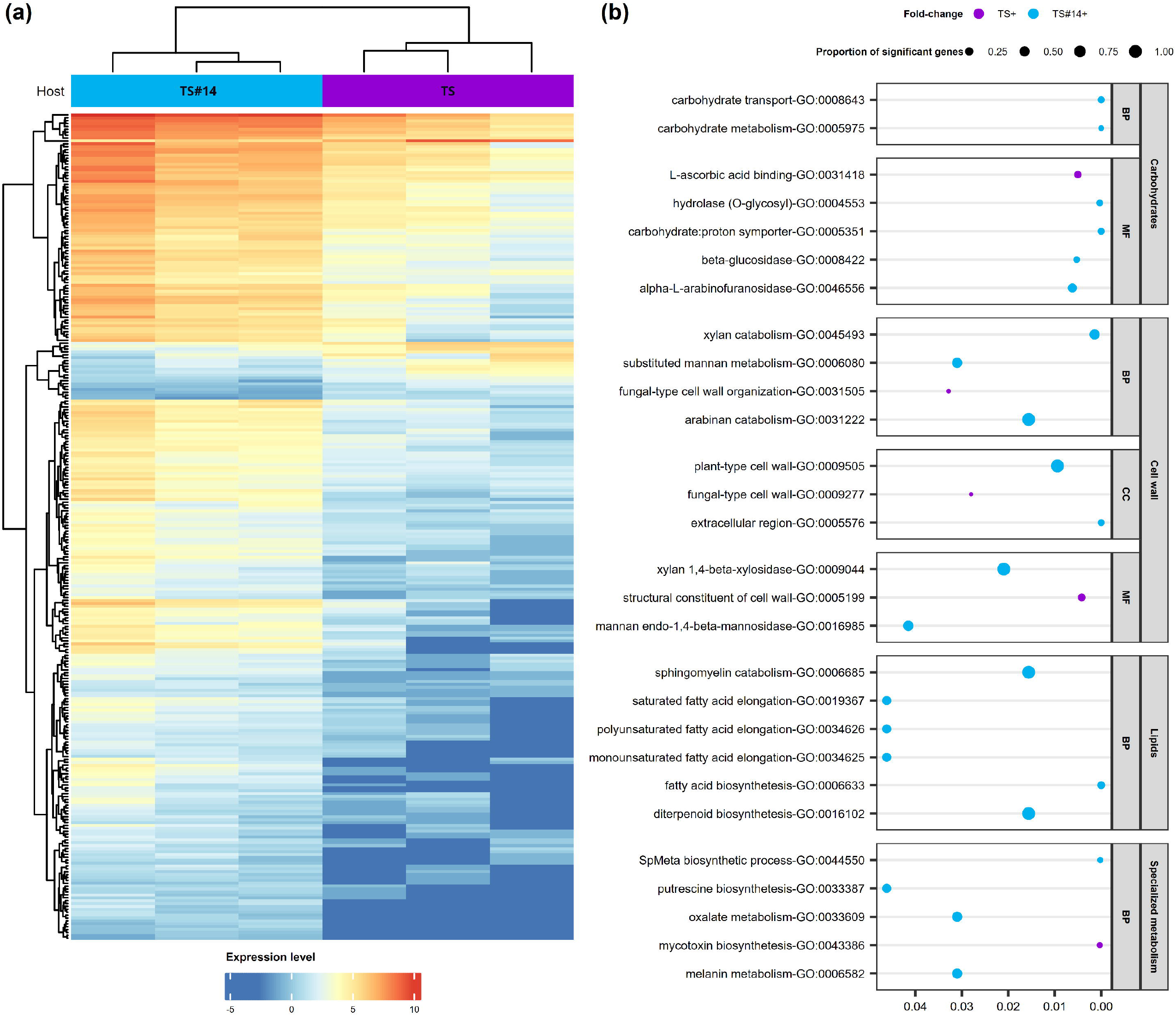
Global transcriptional response of *Botrytis cinerea* to TS and TS#14 *Vitis* lines. (a) Heatmap of differentially expressed *Botrytis* genes (DEGs) across infected TS (purple) and TS#14 (blue) plants. (b) Gene Ontology enrichment analysis of *Botrytis* DEGs. *B: biological process, C: cellular compartment, M: molecular function. Blue: GO term significantly over-represented in genes induced when B. cinerea grows on TS#14; Purple: GO term significantly over-represented in genes induced when B. cinerea grows on TS*.

GO enrichment analysis further highlighted distinct strategies depending on the host background (Supplementary Table S6). On TS wild-type, enriched terms were associated with cell wall reorganization, iron and copper ion transport, and the synthesis of toxins, pointing to a stress-related response aimed at maintaining viability and weakening host defenses (Figure 5b). Copper/iron homeostasis and detoxification are hallmarks of fungal adaptation to host-derived oxidative stress and contribute to *B. cinerea* fitness and pathogenicity (Robinson et al., 2021; Rodríguez-Ramos et al., 2023), while secondary metabolite pathway (*e*.*g*., botrydial / botcinic acid) underpin virulence under restrictive conditions (Cheung et al., 2020).

In contrast, infection on TS#14 induced a much broader set of functions, including plant cell wall polysaccharide (xylans, mannans, arabinans) catabolism, sugar transport, and the metabolism of lipids and specialized compounds such as putrescine. Upregulation of hemicellulose-targeting CAZymes together with monosaccharides transporters is a recurrent signature of successful colonization, enabling efficient uptake of host-derived sugars (Blanco-Ulate et al., 2014; Srivastava et al., 2020; Van Kan et al., 2017). Moreover, activation of polyamine (putrescine) metabolism is compatible with roles of fungal polyamines in growth and stress tolerance. In *B. cinerea*, inhibition of ornithine/arginine decarboxylases impedes growth and can be rescued by exogenous putrescine (Smith et al., 1990; Valdés-Santiago and Ruiz-Herrera, 2014).

Altogether, these results suggest that *B. cinerea* faces a more hostile environment growing on TS wild-type host and primarily deploys survival and virulence strategies, whereas growing on TS#14 host it encounters favorable conditions that allow direct exploitation of host-derived nutrients. In other words, while the pathogen struggles to establish itself on TS wild-type, it appears to engage a broad nutrient-acquisition program on TS#14. Mechanistically, our radiotracer data show that overexpression of VvHT5 increases active glucose uptake without altering bulk soluble sugar pools in leaves (Supplementary Figure S1e), indicating a shift in flux and local sink activity rather than a rise in steady-state sugar levels. This is consistent with the view that sink strength and apoplastic sugar turnover, coordinated by transport processes and extracellular sucrose cleavage, can change markedly while pool sizes remain buffered (Roitsch and González, 2004; Ruan, 2014). In this light, *B. cinerea* transcriptome on TS#14, enriched for hemicellulose-targeting CAZymes and sugar transporter, fits a nutrient-acquisition program engaged under dynamic host-carbon supply, a pattern widely observed during successful colonization across hosts (Blanco-Ulate et al., 2014; Liu et al., 2022; Ma et al., 2022)(Blanco-Ulate et al, 2014; Ma et al, 2022; Liu et al, 2022).

### 3.8. Sugar transporters in *Vitis* and *Botrytis* during infection

Because the functional enrichment analysis in *B. cinerea* highlighted sugar transport as a major process associated with infection on TS#14 and given that *VvHT5* itself encodes a sugar transporter, we next examined the transcriptional regulation of sugar transporter families in both host and pathogen.

In grapevine (Figure 6a), *VvHT5* reached similar expression levels in both genotypes at 48 hpi. Two other transporters, *INT2* and *CWINV5*, clustered with *VvHT5*, indicating that they follow a similar expression pattern during infection. Several transporters, including *ERD6-like16, HT4* and *HT13*, were expressed at a basal level and further induced upon infection, consistent with their involvement in pathogen-triggered sugar mobilization, suggesting that they may participate in the host transcriptional reprogramming associated with sugar partitioning. By contrast, *ERD6-like4, SWEET4* and *CWINV1*, were undetectable under both non-treated and mock conditions but strongly induced during infection, pointing to a pathogen-specific activation. The co-induction of cell-wall invertase and *SWEET4* is consistent with an apoplast-facing mobilization module that increases local hexose supply and sugar efflux at infection sites. Notably, *VvSWEET4* is robustly upregulated by *B. cinerea* and contributes to the interaction in grapevine (Chong et al., 2014; Meteier et al., 2019). Finally, some transporters such as *SWEET10* and *TMT3* were downregulated upon infection. This aligns with a shift away from apoplastic efflux and vacuolar storage toward immediate cytosolic utilization during stress, as expected for tonoplast monosaccharide transporters and consistent with system-level views of sugar transport and signaling in grapevine (Lecourieux et al., 2014; Wormit et al., 2006). Strikingly, none of these transporters seemed to show differential expression between TS#14 and TS wild-type, underlining the absence of functional redundancy that could buffer the effect of *VvHT5* overexpression. Accordingly, divergence between genotypes at 48 hpi likely resides at the level of flux control and post-translational regulation rather than steady-state mRNA abundance, a common feature for sugar carriers under pathogen pressure (Lecourieux et al., 2014; Lemonnier et al., 2014). In addition, sugar transporter expression pattern is consonant with Monnereau et al (2025), who documented a coactivation of *VvHT5* and cell-wall invertase during *B. cinerea* challenge together with a contribution of SWEET transporters to infection-induced carbon partitioning in leaves. This reinforces the convergence of infection signature across systems. In this study, *A. thaliana* ectopically overexpressing *VvHT5* exhibited increased active glucose uptake, providing mechanistic support for a flux-driven, apoplast-oriented response.

**Figure 6.**
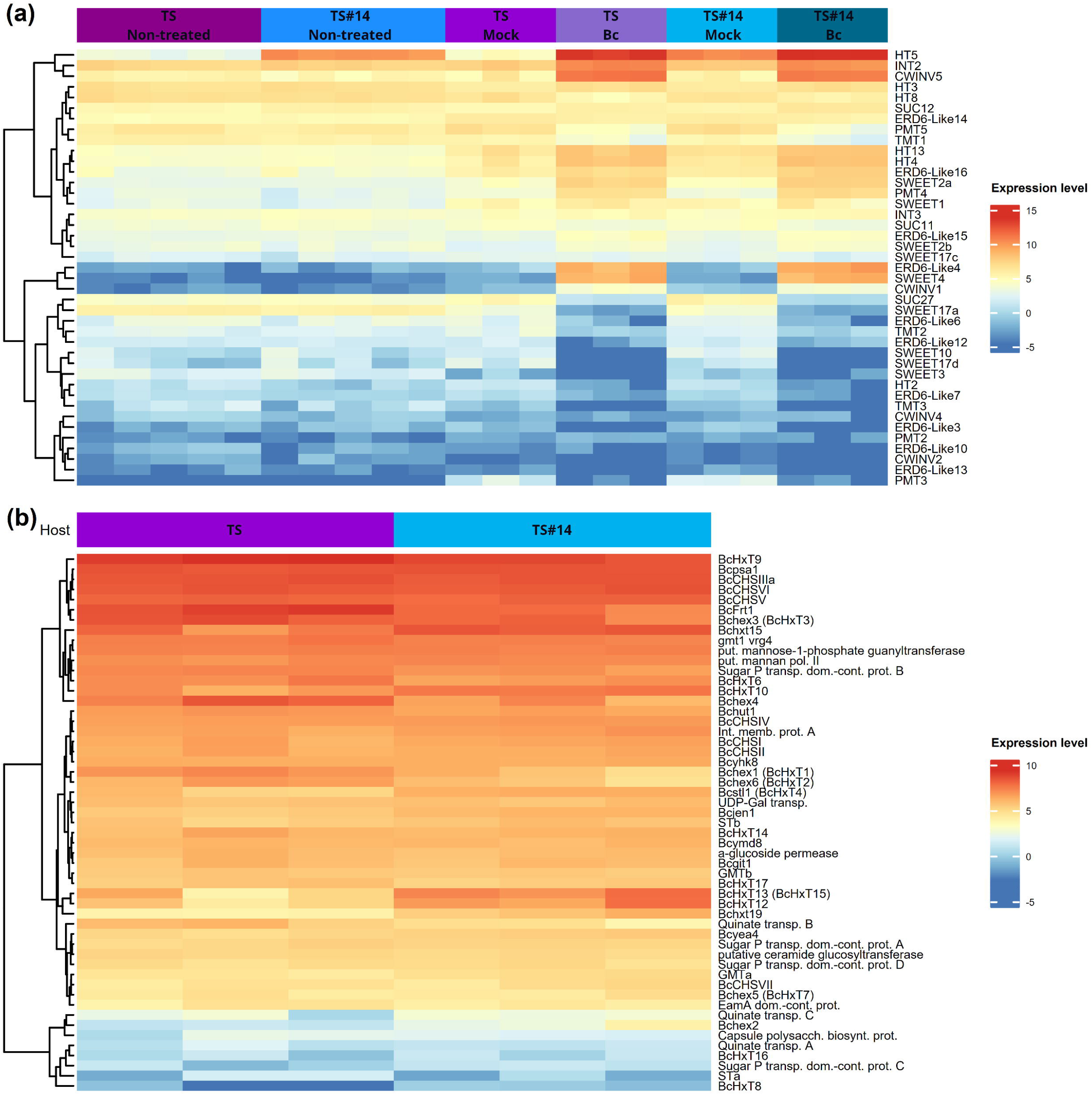
Differential regulation of sugar transporters during the *Vitis–Botrytis* interaction. (a) Expression profile of sugar transporter genes in *Vitis vinifera. Dark purple: TS Non-treated, Blue: TS#14 Non-treated, Purple: TS Mock-inoculated, Light purple: TS Bc-infected, Light blue: TS#14 Mock-inoculated, Dark blue: TS#14 Bc-infected*. (c) Expression profile of sugar transporter genes in *Botrytis cinerea. Purple: TS, Blue: TS#14*.

In *B. cinerea* (Figure 6b), sugar transporter expression revealed a striking asymmetry between the two host lines. A large set of hexose transporter (*BcHxT8, BcHxT10, BcHxT12, BcHxT13, BcHxT15, BcHxT17, BcHxT19*) were preferentially expressed during infection of TS#14 compared to TS wild-type, while others (*BcFRT1, Bchex1, Bchex3, Bchex4, Bchex6, BcHxT4, BcHxT6, BcHxT9*) were preferentially expressed during infection on TS wild-type. This pattern highlights functional specialization within the *B. cinerea* transporter repertoire, depending on the host background. Rather than indicating a simple increase in sugar availability in TS#14, these data suggest that the altered sugar dynamics resulting from *VvHT5* overexpression shape the pathogen’s transcriptional program, leading to the activation of distinct subsets of fungal transporters adapted to each host environment. Consistent with this view, *B. cinerea* encodes a multi-gene sugar-uptake system that includes the high-affinity fructose/H^+^ symporter BcFRT1 and an expanded BcHxT family whose members differ in substrate preference and induction regimes (Doehlemann et al., 2005; Dulermo et al., 2009). Functional genetics further supports specialization within this repertoire. BcHXT15 and context-dependently BcHXT19 contribute to D-galacturonic acid (GalA), the principal monomer of pectin, uptake downstream of pectin depolymerization (Zhang et al, 2011; Zhang & van Kan, 2012). Consistently, GalA constitutes an important carbon source for *B. cinerea*. Mutants defective in GalA catabolism display impaired growth on GalA and attenuated virulence on *A. thaliana* and *Nicotiana benthamiana*, underscoring the contribution of GalA utilization to *in planta* fitness (Zhang et al., 2011; Zhang and van Kan, 2013). Upstream regulation by BcGaaR coordinates galacturonic acid/pectin utilization and biases transporter subset according to cell wall-derived sugars and local conditions (Zhang et al., 2016). Given the over-representation of *B. cinerea* hemicellulose-depolymerizing CAZymes expressed on TS#14 host, it is highly likely that hemicellulose breakdown is increased, thereby releasing more pentoses (D-xylose, L-arabinose) and hexoses (*e*.*g*., mannose, galactose), implying a requirement for coordinated uptake. While direct substrate assignments remain limited for many *B. cinerea* Major Facilitator subfamily (MFS) carriers, infection transcriptomes repeatedly show concerted induction of transporter genes alongside CAZymes, consistent with a coupled “depolymerize-and-import strategy” (Blanco-Ulate et al., 2014; Dulermo et al., 2009; Reboledo et al., 2021). It is noteworthy that in filamentous fungi, transporter expression and uptake capacity correlate for pentoses and GalA (Tamayo et al., 2024).

Practically, these data argue that *VvHT5* reshapes host carbon dynamics rather than simply raising hexose levels, thereby selecting distinct transporter subsets in the fungus. Within this framework, the TS#14-biased vs TS wild-type subset of hexose transporters likely reflects the environment-dependent partitioning of the BcHxT repertoire rather than a uniforme rise in sugar supply. Multigene HXT systems in *B. cinerea* exhibit distinct induction profiles across carbon sources and pH, enabling flexible assimilation, and their deployment is further modulated by carbon catabolite control (Dulermo et al., 2009; Wu et al., 2020; Zhang et al., 2014).

## 4. CONCLUSION

Our study demonstrates that overexpression of the sugar transporter VvHT5 in grapevine fundamentally alters host–pathogen interactions with *Botrytis cinerea*. Transgenic lines with enhanced *VvHT5* expression exhibit increased active glucose uptake without modifying bulk soluble sugar pools, indicating a shift in sugar flux rather than steady-state levels. This altered carbon dynamics correlates with heightened susceptibility to *B. cinerea*, particularly in the more susceptible Thompson Seedless cultivar, accompanied by larger necrotic lesions and higher fungal biomass. Transcriptomic analyses reveal that *VvHT5* overexpression has minimal constitutive effects but triggers profound host- and pathogen-specific transcriptional reprogramming during infection. Grapevine defense pathways, including jasmonic acid signaling, cell wall remodeling, and specialized metabolism, are misregulated in TS#14, while *B. cinerea* responds by upregulating sugar transporters, CAZymes, and nutrient acquisition programs. Notably, VvHT5 and its Arabidopsis ortholog AtSTP13 are both induced by *B. cinerea*, yet their outcomes differ: VvHT5 appears exploited by the pathogen, increasing susceptibility, whereas AtSTP13 induction in Arabidopsis limits apoplastic sugar, enhancing tolerance. This contrast underscores that insights from model plants may not fully predict crop responses. These findings establish STP13-like transporters as pivotal regulators of sugar-mediated host–pathogen interactions and highlight the species-specific interplay between sugar flux, plant defense, and pathogen exploitation.

## AUTHOR CONTRIBUTION

B.M., O.Z., P.C.T., S.L.C. – Conceptualization, Methodology, Project administration, Supervision, Funding acquisition

B.M., C.G., V.L., P.V. – Investigation, Data curation

B.M., C.C., C.G., S.L.C. – Formal analysis, Validation, Visualization

B.M., C.C., S.L.C. – Writing – original draft, Writing – review & editing

All authors – Read and approved the final manuscript

## ACKNOWLEDGMENTS

This work was supported by the Centre National de la Recherche Scientifique, the University of Poitiers, the Region Nouvelle-Aquitaine (BM PhD grant - CryptoCARBO project-AAPR2021-2020-11949210), France Relance program, the State-Region Planning Contracts (CPER) and the European Regional Development Fund (FEDER). This work has benefited from the facilities and expertise of ImageUP platform (UAR CNRS 2038—University of Poitiers). We are grateful to Katerina Labonova, whose expertise and involvement in the grapevine transformation procedures were essential for generating the transgenic material used in this study We thank all colleagues and technical staff for inspiring discussions.

## DATA AVAILABILITY

RNA-seq raw reads have been deposited in the NCBI Sequence Read Archive under the BioProject accession number PRJNA1363860.

## SUPPLEMENTAL DATA

**Supplementary Figure S1. Additional characterization of *VvHT5-*overexpressing *Vitis* lines**. (a) Greenhouse-acclimated TS WT and TS#14 plants. (b) *GFP* expression levels in leaves of wild-type and transgenic TS and Cha lines. Gene expression was quantified by qRT-PCR from healthy mature leaves and normalized to the reference gene *VvGAPDH*. Data represent mean ± SD (three biological replicates). (b) Accumulation of the GFP–VvHT5 protein in transgenic TS and Cha lines assessed by Western blot. Microsomal fractions extracted from healthy mature leaves were probed with anti-GFP antibodies. Black arrows indicate GFP-specific signals. (d) Detection of GFP signal in TS transgenic line TS#14 overexpressing GFP–VvHT5. Confocal images of leaf epidermis are shown alongside TS WT as control. GFP fluorescence is shown in green and chlorophyll autofluorescence in red. Scale bars = 10 µm. (e) Soluble sugar content in leaves of *VvHT5*-overexpressing *Vitis* lines. Soluble sugars were quantified from whole healthy leaves. Green, blue, and yellow bars represent mean sucrose, glucose, and fructose contents, respectively; black dots represent individual measurements (five biological replicates for TS lines and two for Cha lines).

**Supplementary Table S1**. List of primer sequences for PCR amplifications

**Supplementary Table S2**. Raw read count matrix for all samples used in RNA-seq analyses. *Values correspond to unnormalized gene-level counts for both Vitis vinifera and Botrytis cinerea, prior to differential expression analysis*

**Supplementary Table S3**. Statistical analysis of total, active and passive (CCCP-insensitive) glucose uptake in leaves of wild-type and *VvHT5*-overexpressing transgenic lines, according to pairwise Wilcoxon test (alternative = “less”) using Benjamini-Hochberg adjustment method (\**p-*value < 0.05, \*\**p-*value < 0.01, \*\*\**p-*value < 0.001).

**Supplementary Table S4**. Statistical analysis of lesion size distribution from leaf discs of wild-type and VvHT5-overexpressing transgenic lines inoculated with *B. cinerea*, according to pairwise Chi square test using Benjamini-Hochberg adjustment method (\**p-*value < 0.05, \*\**p-*value < 0.01, \*\*\**p-*value < 0.001)

**Supplementary Table S5**. Enriched GO term in *Vitis vinifera* cv Thompson Seedless wild-type and VvHT5-overexpressing line TS#14 under *B. cinerea* pressure (48 hpi)

**Supplementary Table S6**. Enriched GO term in *B. cinerea* growing on *V. vinifera* cv Thompson Seedless wild-type orVvHT5-overexpressing line TS#14 (48 hpi)

