## Supplementary figures and images for "Overexpression of the sugar transporter *VvHT5* turns grapevine into a better host for *Botrytis cinerea*"

### Supplementary Figure S1

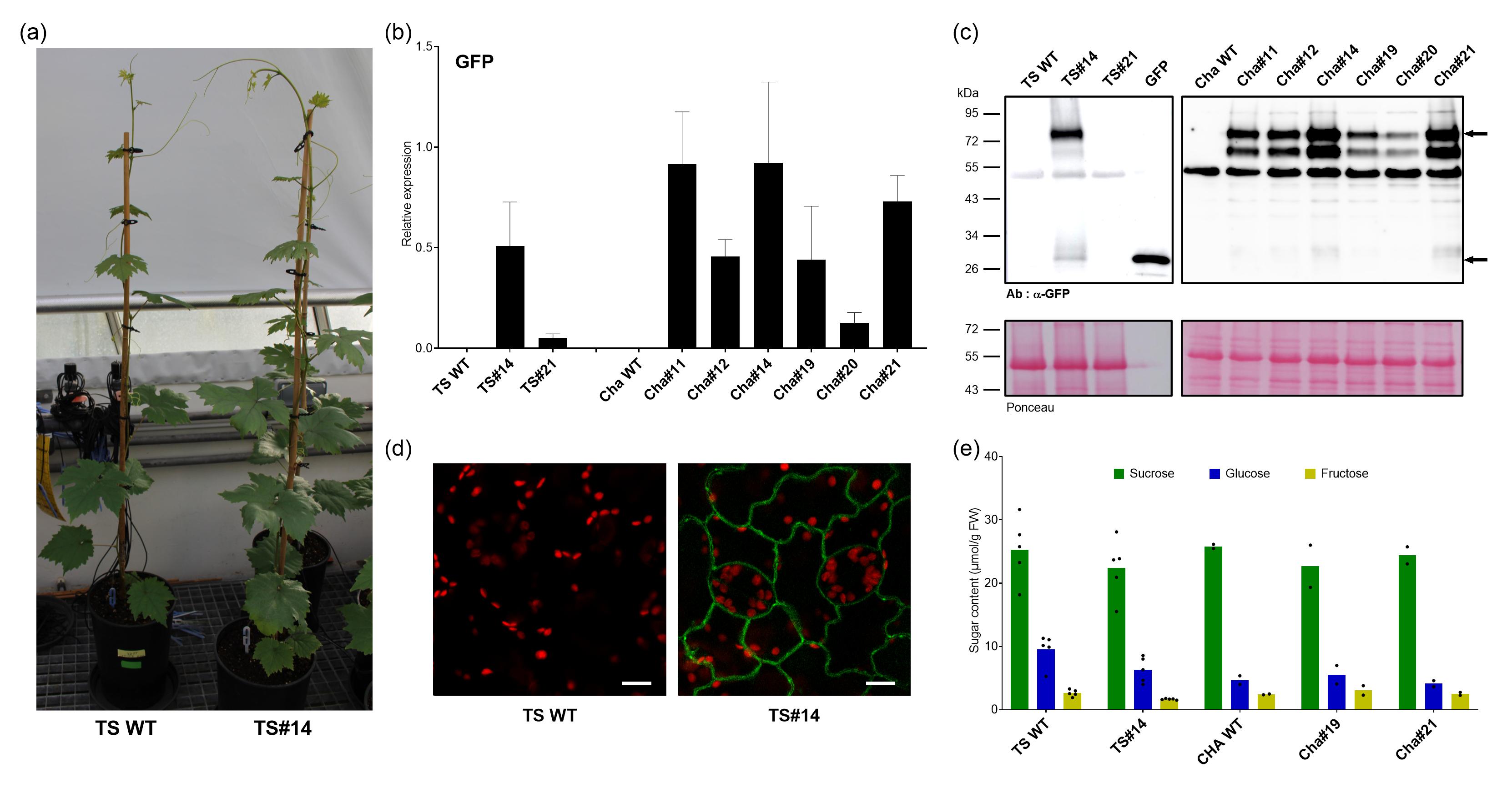
